# Iron nanoparticle-labeled murine mesenchymal stromal cells in an osteoarthritic model persists and demonstrates anti-inflammatory mechanism of action

**DOI:** 10.1101/571976

**Authors:** Amanda M. Hamilton, Wing-Yee Cheung, Alejandro Gómez-Aristizábal, Sayaka Nakamura, Amélie Chaboureau, Shashank Bhatt, Anirudh Sharma, Mohit Kapoor, Paula J. Foster, Sowmya Viswanathan

**Affiliations:** Imaging Research Laboratories, Robarts Research Institute, London, ON, Canada; The Arthritis Program, Toronto Western Hospital, Toronto, ON, Canada; Division of Genetics and Development, Krembil Research Institute, University Health Network, Toronto, ON, Canada; Department of Surgery, Department of Laboratory Medicine and Pathobiology, University of Toronto, Toronto, ON, Canada; Department of Medical Biophysics, Schulich School of Medicine and Dentistry, Western University, London, ON, Canada; Institute of Biomaterials and Biomedical Engineering, University of Toronto, Toronto, ON, Canada; Department of Medicine, University of Toronto, Toronto, ON, Canada

## Abstract

Osteoarthritis (OA) is characterized by cartilage degradation and chronic joint inflammation. Mesenchymal stem cells (MSCs) have shown promising results in OA, but their mechanism of action is not fully understood. We hypothesize that MSCs polarize macrophages, which are strongly associated with joint inflammation to more homeostatic sub-types. We tracked ferumoxytol (Feraheme™, iron oxide nanoparticle)-labeled murine MSCs (Fe-MSCs) in murine OA joints, and quantified changes to homeostatic macrophages.

10-week-old C57BL/6 male mice (n=5/group) were induced to undergo osteoarthritis by destabilization of medical meniscus (DMM) or sham surgery. 3 weeks post-surgery, mice were injected intra-articularly with either fluorescent dye-(DiR) labeled or DiR+ferumoxytol-labeled (DiR+Fe) bone marrow mesenchymal stem cells (MSC, 50×10^3^ MSCs/mouse) or saline (control), to yield 4 groups (n=5 per group for each timepoint [1, 2 and 4weeks]): i) DMM+Saline; ii) DMM+DiR+Fe-MSC; iii) DMM+DiR MSC; iv) SHAM+DiR+Fe-MSC and saline in contralateral knee. 4 weeks after injection, mice were imaged by MRI, and scored for i) OARSI to determine cartilage damage and ii) immunohistochemical changes in CD206, F480 and Prussian Blue/Sca-1 to detect homeostatic macrophages, total macrophages and ferumoxytol-labeled MSCs respectively.

Ferumoxytol-labeled MSCs persisted in DMM knee joints at greater levels than in SHAM-MSC knee joints. We observed no difference in OARSI scores between MSC and vehicle groups. Sca-1 and Prussian Blue co-staining confirmed the ferumoxytol label resides in MSCs, although some ferumoxytol label was detected in proximity to MSCs in macrophages, likely due to phagocytosis of apoptotic MSCs, increasing functionality of these macrophages through MSC efferocytosis. We showed decreased MRI synovitis scores in MSC-treated compared to control animals. For the first time we show that MSC-treated OA mice had increased macrophage infiltration (p<0.08) with an increased proportion of CD206+ (homeostatic) macrophages (p<0.01), supporting our hypothesis that MSCs modulate synovial inflammation.

## Introduction

Osteoarthritis (OA) is a common joint disease affecting 1 in 10 Canadians and is expected to increase to 1 in 4 by 2040. It is a lasting condition in which cartilage breaks down, causing bones to rub against each other, resulting in stiffness, pain and loss of joint movement. Currently, there are few effective treatments available for patients suffering from osteoarthritis. **M**esenchymal **S**tromal **C**ells (MSCs) are cells that can be obtained from bone marrow and other tissues. Early clinical data shows improvements in cartilage volume and quality [1–6] and secretion of immunomodulatory factors. Our pioneering Canadian trial (NCT02351011, clinicaltrials.gov) using autologous bone-marrow-derived MSCs showed significant improvement in symptoms and quality of life relative to baseline in osteoarthritic patients; for the first time, we showed clinically that MSCs reduced synovial joint levels of pro-inflammatory macrophages, a potent inflammatory mediator, suggestive of a possible mechanism of action [7]. Macrophages are the most prevalent immune cell in OA joints [8], are elevated in OA compared to healthy joints [9], and contribute to synovial inflammation and fibrosis, characteristic of OA.

Understanding the immediate post-transplant joint environment by visualizing and monitoring localization and persistence of MSCs in a direct, non-invasive, longitudinal manner will address a major barrier in evaluating the success of MSC therapies in OA. Although early clinical trial results with MSCs are promising, unanswered questions regarding dosing, timing, frequency, interaction with macrophages, and mechanism of action of MSC persist.

Previously MSCs have been imaged in animal models using superparamagnetic iron oxide nanoparticles (SPIO) such as such as ferumoxides and ferucarbotran (i.e. Feridex™, Endorem^®^ and Resovist™) because of their ease of use in stem cells, greater sensitivity for MR imaging compared to other contrast agents including gadolinium and CellSense, and lack of requirement for specialized multinuclear MRI systems for visualization. Ultrasmall SPIOs (i.e. ferumoxytol or Feraheme™) are introduced into cells by endocytosis and require transfection protocols using protamine sulfate and heparin [10]. Ferumoxytol has been clinically-approved by Health Canada and the Food and Drug Administration (FDA) for treatment of anemia in chronic-kidney disease patients, making it an attractive cell tracking agent.

Adipose-tissue MSCs have previously been labeled with ferumoxytol and feasibility of imaging *in vitro* and *in vivo* in rats models of osteochondral defects has been shown [11]. Other studies have also imaged MSCs labeled with other iron-nanoparticles and demonstrated persistence out to 5 weeks [12] in Hartley strain guinea pigs, and persistence out to 6 months in immunosuppressed NOD-SCID mice [13].

In this study, we attempted to demonstrate feasibility of labeling and MRI tracking of murine MSCs in a murine joint model of mildly inflammatory osteoarthritis that is clinically translatable and relevant. This study serves as a pre-requisite for subsequent clinical translation involving ferumoxytol-labeling of human MSCs intra-articularly injected into knee OA patients. Importantly, we wanted to identify a putative anti-inflammatory mechanism of action of MSCs through polarization of monocytes/macrophages in the arthritic joint. Monocytes/macrophages are the most prominent immune effects in OA and their levels correlate with OA severity, which we have recently also correlated with patient-reported pain and symptoms [14].

## Materials & Methods

### Cell isolation

Mouse bone marrow stromal mesenchymal cells were isolated from mouse long bones. Briefly, bone marrow was flushed out of mouse long bones (mouse tibiae, femurs), and were cultured in low glucose DMEM, supplemented with 10% FBS and Glutamine for less than 14 days.

### Animals

10 week-old C57BL/6 male mice (n=5/group, Jackson Laboratories) were used for our studies. Mice underwent surgery (destabilization of medial meniscus (DMM) or sham surgery) and injected with either fluorescent dye-(DiR) labeled or DiR+ferumoxytol-labeled (DiR+Fe) mesenchymal stem cells (MSC, 50×10^3^ MSCs/mouse) or saline (control), and were thus separated into 4 treatment groups: i) DMM+Saline; ii) DMM+DiR+Fe-MSC; iii) DMM+DiR MSC; iv) SHAM+DiR+Fe-MSC and saline in contralateral knee (Fig 1). All mice were fed ad libitum and were allowed to roam freely in cages throughout the study.

**Figure 1:**
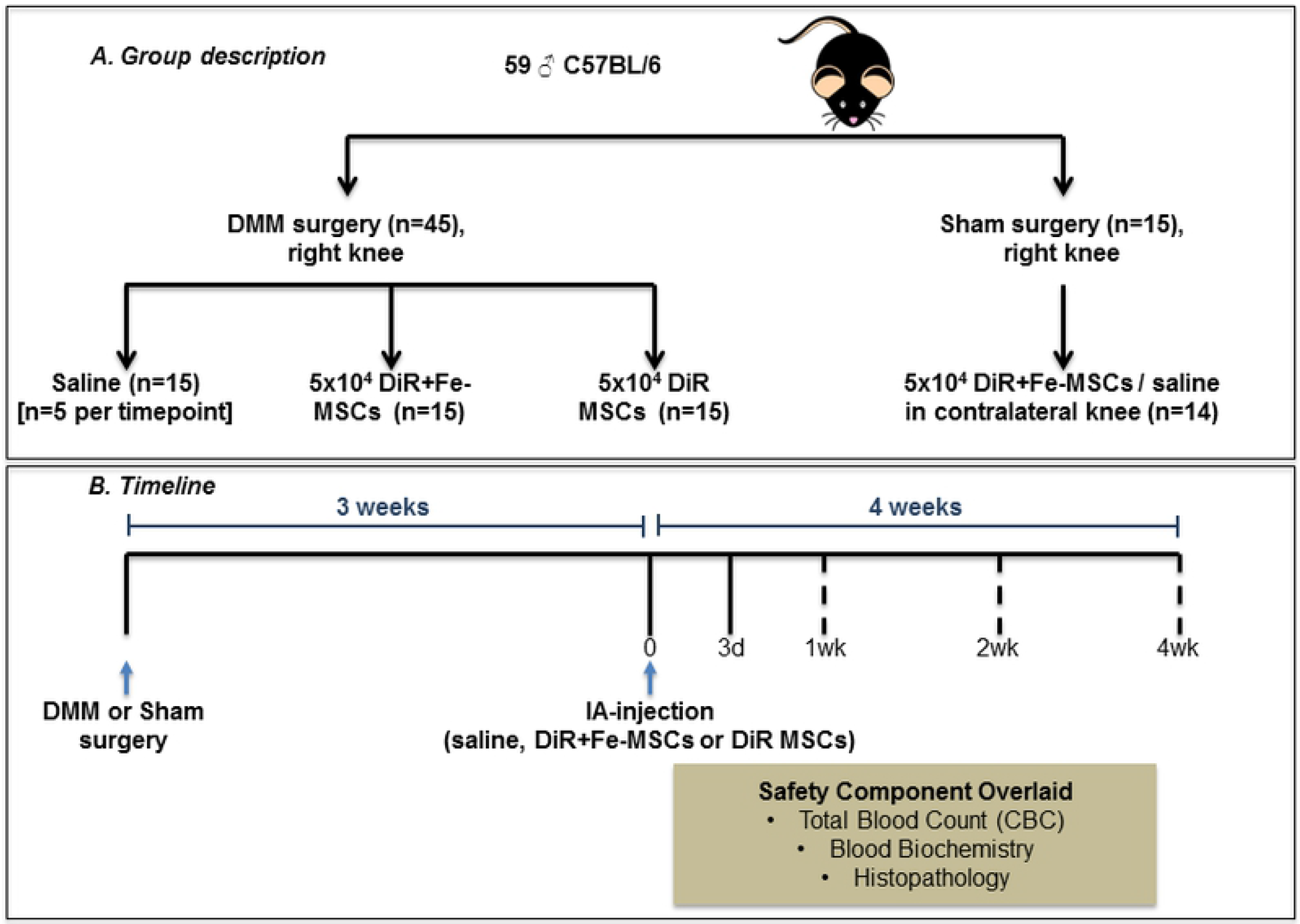
Experimental schematic of the DMM model. **A.** Group description: 10 week old male C57BL/6 mice were subjected to DMM or SHAM surgery. 3 weeks post-DMM surgery, mice were injected intra-articularly with saline, DiR+ferumoxytol-labeled MSCs (DiR+Fe-MSCs), or only DiR-labeled MSCs (DiR-MSCs) in the right knees (n=5/timepoint). SHAM operated mice were injected with MSCs in right knees, and saline on left knees. (n=4 for 1 week, n=5 for 2 and 4 week timepoints). DiR: 1,1’-Dioctadecyl-3,3,3’,3’-Tetramethylindotricarbocyanine Iodide. **B.** Timeline: Mice were assessed for total blood count, blood biochemistry, histopathology and MSC persistence in knee joints via MRI (On day 0 and day 3, and 1, 2 or 4 weeks post-injection). Dotted lines indicate that only one of the timepoints were selected for MRI.

### Surgical induction of instability

DMM – DMM surgery was performed as described by Glasson et al. [15]. Briefly, mice were induced with 2% isoflurane, and their right knees were shaved and prepped with iodine. At 90 degrees of flexion, the joint line was identified using a straight sharp tip micro probe (0.025 mm) facilitating the establishment of a medial para-patellar intra-articular port. A 3 mm skin incision was made over the medial portal using a knife with a number 10 blade. Following this, the capsule was incised in line with the skin incision medial to the patellar ligament. With the help of a microcurette (0.5 mm), the fat pad was dissected and the medial meniscotibial ligament (MMTL) was exposed. The MMTL is a translucent transverse band that connects the medial meniscus (MM) to the tibial plateau. Under direct vision, the MMTL was incised using a straight tip vannas spring scissors (0.075 mm). Care was taken not to damage the cartilage or other soft tissue structures. During the destabilization, bleeding encountered was controlled with direct pressure using absorption spears. The destabilization was confirmed using a probe against the freely mobile medial meniscus. The capsule and skin were then sutured (8-0). Buprenorphine (0.2 mg/mL, Indivior) was injected subcutaneously at the time of surgery and 24 hours post-surgery for pain maintenance. Mice were allowed to roam freely in their cages after surgery, and monitored closely for post-surgical complications.

### Cell labeling and injection

A subset of research animals (DMM and sham) received ferumoxytol-labeled MSC. Prior to intra-articular injection, MSCs were incubated for 2h with 50 μg Fe/mL ferumoxytol (Feraheme™; AMAG Pharmaceuticals Inc., Waltham, MA, USA) as per Thu et al in serum-free DMEM containing 3 U/mL heparin and 60 mg/mL protamine sulfate at 37°C, 5% CO_2_ [10]. After this initial incubation, cells were supplemented with complete low glucose DMEM (containing 10% FBS) and further incubated at 37°C, 5% CO_2_ overnight. After labeling, cells underwent sequential PBS washing before and after dissociation. All cells (Fe-MSCs and unlabeled MSCs) were stained with the lipophilic fluorescent dye DiR (also called DiIC18(7), full chemical name: 1,1’-Dioctadecyl-3,3,3’,3’-Tetramethylindotricarbocyanine Iodide) for 30 minutes and then washed three times with PBS before quantification. DiR is a lipophilic, near-infrared fluorescent cyanine dye. Cell viability was assessed by trypan blue exclusion assay, and cells were subsequently intra-articularly injected into mice in accordance to their respective groups (Fig 1).

### Intra-articular knee injections

Mice were induced with 2% isoflurane, and their knees were depilated, prepped and positioned as in the DMM surgery. The patellar ligament was identified as a pearl-white vertical band. Using a 10 μL Hamilton Syringe with custom 30G needle, the joint line was identified, and the needle was inserted along the medial border of the patellar ligament, keeping it in line with the joint line, slowly advancing it till loss of resistance was appreciated. Following this, the plunger was slowly pushed to deliver 4 μL of DiR+Fe-MSCs or DiR MSCs (5×10^4^ cells) into the joint space. Once the MSCs were delivered, the needle was removed, and the knee joint was flexed and extended slowly. Saline was used as a vehicle control treatment.

### Magnetic resonance imaging and analysis

All MRI examinations were performed using a 3 Tesla (T) GE Discovery MR750 whole-body clinical MR scanner with a custom-build high-performance gradient insert coil (maximum gradient strength = 500 mT/m and peak slew rate = 3000 T/m/s) and a custom solenoid radiofrequency mouse body coil (4 cm long, 3 cm diameter). Mice were anaesthetized with 2% isoflurane. Whole-body mouse images were acquired with the following three dimensional balance steady state free precession (3D bSSFP) imaging parameters: 200 mm isotropic spatial resolution, with a 5-cm field of view (FOV) and 250 x 250 image matrix and a 200 μm slide thickness; phase FOV = 0.5; repetition time/echo time (TR/TE): 6.3/3.152; flip angle=35°; bandwidth=±31.25kHz; eight phase cycles; number of excitations=2; scan time=34min. Mice were imaged three times during the course of the experiment: on the day of cell delivery (day 0), 3 days post- and either 1, 2 or 4 weeks post-cell delivery. Images were viewed and analyzed using the open-source DICOM viewer OsiriX (Pixmeo, Switzerland). Void volumes were measured via manual hand tracing and volume calculation algorithms (Osirix) to assess presence of MSCs in joint space over time. MR images of DMM surgical knees were also scored for suspected evidence of inflammation. Scoring was conducted on a scale of 1 to 5 where 1 indicated no evidence of inflammation (resemblance to control non-surgical knees) and 5 indicated clear evidence of inflammation including increased total knee volume and hyperintensity of tissue surrounding the knee joint.

### Viability

Viability of unlabeled and ferumoxytol-labeled murine MSCs (n=4) and human MSCs (n=3) was assessed by Trypan blue dye exclusion using ViCell (Beckman Coulter).

### Tri-lineage differentiation

Unlabeled and ferumoxytol-labeled murine MSCs were tested for chondrogeneic and osteogeneic differentiation potential using MesenCult (StemCell Technologies) differentiation as per manufacturer’s instructions. Unlabeled and ferumoxytol-labeled murine MSCs were tested for adipogeneic differentiation potential using StemPro differentiation kit (Life technologies) or using low glucose DMEM media (Life technologies) supplemented with 10% FBS (Fisher), 1% Glutamine (Life technologies), 1 μM dexamethasone (Sigma), 5 μg/mL insulin (Sigma), 0.125 mM indomethacin (Sigma) and 0.5 mM 3-Isobutyl-1-methylxanthine (Sigma). Unlabeled and ferumoxytol-labeled human MSCs were tested for tri-lineage differentiation potential using StemPro differentiation kits (Life technologies) for adipogeneic, chondrogeneic and osteogeneic differentiation as per manufacturer’s instructions. Staining for murine and human MSCs was performed using Oil Red O (Sigma) stain for adipogenesis, Alcian Blue 8GX (Sigma) for chondrogenesis and Alizarin Red S (Sigma) for osteogenesis.

### Gene expression

Unlabeled and ferumoxytol-labeled murine MSCs were stimulated with 30 ng/mL murine interferon-gamma (Peprotech) at 37°C, 5% CO2 for 18-20 hours. Post-stimulation, total RNA was isolated using TRIzol^®^ Reagent (Life Technologies) and converted into cDNA with High-Capacity cDNA RT Kit (Life Technologies). Real-time RT PCR was performed using FastStart Universal SYBR Green Master mix (Roche). Samples were analyzed in triplicate for β2-Microglobulin (B2M), inducible NO synthase (iNOS), Interleukin 10 (IL10), Transforming growth factor beta (TGFβ1), Interleukin 6 (IL6), Hepatocyte growth factor (HGF), Programmed death-ligand 1 (PDL-1 or CD74) and Cyclooxygenase 2 (COX2 or Prostaglandin endoperoxide synthase 2, PTGS2) (see Table S1 for detailed primer sequences). Thermal cycling was performed with QuantStudio5 (ThermoFisher): 95°C, 2 min followed by 40 cycles of 95°C, 15s and 60°C, 20s. Relative fold change in expression levels were calculated using the 2^-ΔΔCt method, with β2-Microglobulin (B2M) as housekeeping gene [16]. Comparison was determined using paired Student’s t-test.

**Table S1: Primer sequences used for the PCR**

### Safety analysis

At 2 or 4 weeks after MSC or saline injections, whole blood, serum and heart and spleen tissues for histopathological analysis were collected from DMM and sham mice. Whole blood was collected by cardiac puncture immediately following sacrifice. For complete blood count (CBC) assessment, 100 μL of blood was collected per animal by capillary action into a Microvette^®^ tube (Sarstedt) and stored at 4°C until analysis. To isolate serum, 200 μL of blood was collected by capillary action into a Microvette^®^ tube with serum activators and allowed to clot for 60 minutes. Samples were then centrifuged at 10,000 g for 5 minutes at 4°C and serum was collected into a sterile microcentrifuge tube. Isolated serum was stored at −20 °C until analysis at the Animal Resources Centres of the University Health Network. For histopathological assessment, heart and spleen were collected and fixed in 10% formalin, paraffin embedded, sectioned and stained with standard H&E for gross permit histopathological assessment by trained pathologist at the Pathology Core at The Centre for Phenogenomics.

### Histology

After the final MRI scanning session (at 1, 2 or 4 weeks after MSC or saline injections), mice were sacrificed and knee joints were isolated, fixed in 10% formalin. Mouse joints were decalcified using RDO Rapid Decalcifier (Apex Engineering). Cross sections of 4 mm of the medial aspect of the knee joints were deparaffinized, rehydrated and endogenous tissue peroxidases were quenched using 3% hydrogen peroxide. Sections were then treated with Proteinase K (20 μg/mL) for 15 minutes for antigen retrieval and blocked with DAKO background reducing protein blocking agent (Agilent, X090930-2) prior to 1 hour incubation of CD206 (1:200, R&D systems, AF2535), F480 (1:100, Serotec), Prussian Blue (equal volumes of 20% HCl in dH20 and 10% K_4_Fe(CN)_6_-3H_2_O mixture, Sigma Aldrich respectively) and Sca-1 (1:100, Bio-Rad, MCA2782) at room temperature to detect M2 macrophages, total macrophages, ferumoxytol-labeled MSCs and total cells respectively. Detection was performed using anti-goat secondary antibody (for CD206, 1:1000 Novus Biologicals, NB120-7124), anti-rat secondary antibody (for F480 and Sca-1, 1:500 Novus Biologicals, NBP1-75379) and biotinylated anti-rabbit secondary antibody (for cleaved caspase-3, 1:2000, Vector Labs BA-1000) for 30 minutes, followed by avidin-biotin complex detection system (VECTASTAIN Elite ABC system, Vector Labs PK-6100) for 30 minutes and DAB-horseradish peroxidase substrate detection system. Cells were counterstained with hematoxylin (Vector Labs, H3404) to detect total cells. Sections were subsequently dehydrated and mounted with Permount (ThermoFisher Scientific, SP15-500). Images were taken at 40x and analyzed with Fiji ImageJ software.

### ImmunohistologicalImage analysis

Percentage of positively stained CD206 (homeostatic, “M2-like” cells) or F480 (total macrophages) with respect to total cells (stained with Hematoxylin) was quantified to assess synovial inflammation within the DMM or SHAM-operated knee joints. Briefly, 6-8 images were taken of medial knee joint sections at 40x to encompass the synovial tissue adjacent to the medial meniscus. Fiji (ImageJ) software was used to analyze the images. Specifically, each image underwent Colour Deconvolution to separate brown (DAB) channels and purple (hematoxylin) channels. Subsequent images underwent “Threshold” such that positively stained cells were labeled as black. Positive cells were then quantified using the “Analyze Particles” feature with the following specifications: Size (pixel^2):100-3000 and Circularity (0-0.8), which allowed for the software to provide a cell count for the respective stain. (This semi-automatic method of quantification was verified by doing manual counts of 20 images, which resulted in less than 15% error.)

### Histopathology and OARSI Scoring

Medial knee histological sections were stained with Safranin O dye to assess cartilage degradation and joint inflammation according to the OARSI scoring system by Glasson et al. [17]. Briefly, sections were deparaffinized, rehydrated, and stained with Weigert’s Hematoxylin A and B (ThermoFisher Scientific, 50-317-75 and 50-317-79 respectively, 4 minutes), 0.01% Fast Green (Sigma Aldrich, F7252, 5 minutes), and 0.1% Safranin O (Sigma-Aldrich, S2255, 5 minutes). Sections were then dehydrated and mounted in Permount and imaged at 4x using a brightfield microscope. A score of 0 to 6 was determined for each sample, with 0 indicating lack of cartilage damage, and 6 indicating complete cartilage erosion.

### Synovial Inflammation scoring

Synovial inflammation was determined according to OARSI histopathology methods [18]. Briefly, synovial inflammation is graded from 0-4, where 0 denotes no changes to synovial tissue, and 1-4 denotes the spectrum of synovial inflammation that includes: increased proliferation of subsynovial tissue, increased number of lining cell layers, and an increase in infiltration of inflammatory cells.

## Results

### Viability and phenotypic assessment of ferumoxytol-labeled MSC

There were no differences tri-lineage potential of MSCs labeled with or without ferumoxytol (Fig S1). Trypan blue exclusion assay results demonstrated no difference in the viability of unlabeled and ferumoxytol-labeled murine MSCs, at 98.5±1.9% and 94.3±5.9%, respectively (Fig S2). The fold change in gene expression normalized using β2-Microglobulin (B2M) on a panel of 7 genes (iNOS, IL10, TGFβ1, IL6, HGF, PDL-1, COX2) showed no significant difference between the unlabeled MSC and Fe-MSCs (n=3 per group) as determined by paired Student’s t-test except HGF (*p=0.0341) (Fig S3).

**Figure S1: Tri-lineage potency of ferumoxytol labeled murine and human MSCs are not affected by ferumoxytol. A.** Adipogenesis stained with Oil Red O; **B.** Chondrogenesis stained with Alcian Blue, and **C.** Osteogenesis stained with Alizarin Red S. Scalebar = 25 μm; mMSC: murine MSC; hMSC: human MSC.

**Figure S2: No viability differences in ferumoxytol-labeled murine and human MSCs vs. unlabeled murine and human MSCs.** Viability assessed by Trypan Blue exclusion assay expressed as percentages. **A.** murine MSCs (n=4). **B.** human MSCs (n=3).

**Figure S3: Comparable gene expression between ferumoxytol-labeled murine MSCs and unlabeled murine MSCs.** The fold change in gene expression was normalized using β2-Microglobulin (B2M). None of the genes showed a significant difference between the unlabeled MSC and Fe-MSCs (n=3 per group) as determined by paired Student’s t-test except HGF (*p=0.0341).

### Magnetic Resonance Imaging

Three dimensional whole body anatomical images were acquired that allowed well-defined delineation of experimental mouse knees over time. Representative images of control, sham and DMM surgical knees can be seen in figure 2A. In control mouse knees a hyperintense region corresponding to the fatpad of the tibio-femoral joint was clearly evident (Fig 2A, red arrows). Both sham and DMM surgeries required the dissection of this fatpad to enable access to the joint and the absence of the fatpad was evident in all imaged sham and DMM surgical knees. Mice were imaged on the same day as, at three days post and either 1, 2 or 4 weeks post intra-articular saline or cell delivery. For sham and DMM surgery mice that received Fe-MSC, MR images confirmed the intra-articular localization of the cells by the presence of distinct signal voids in the joint space (Fig 2A, red arrows). These signal voids were analyzed for void volume and fractional signal loss over time (Fig 2B&C).

**Figure 2:**
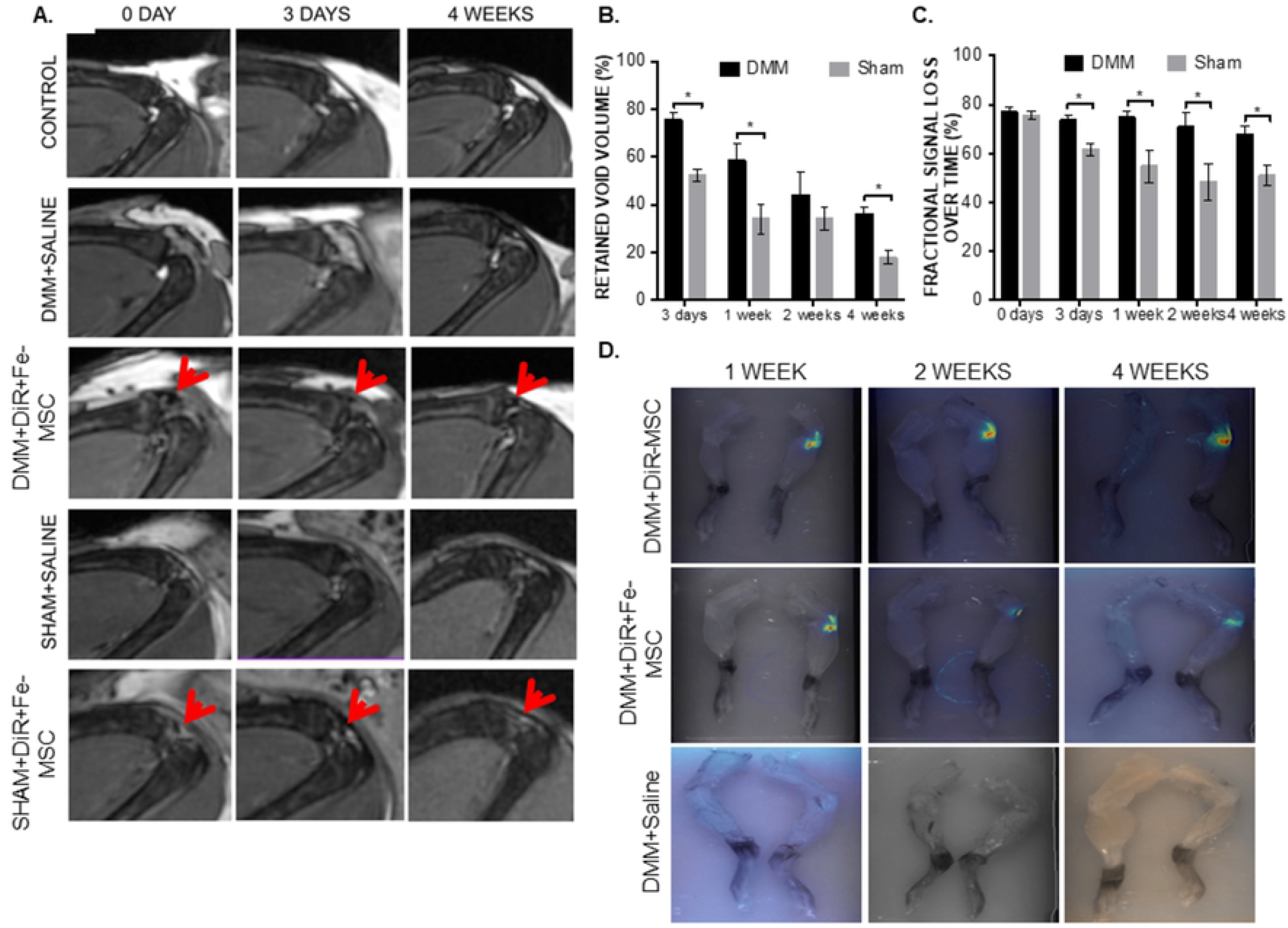
Persistence of MSCs in Knee Joints up to 4 weeks. **A.** All mice received DMM or sham surgery 3 weeks prior to injection with either saline or 5×10^4^ DiR+ferumoxytol-labeled MSCs (DiR+Fe-MSCs) or only DiR-labeled MSCs (DiR MSCs). Mice were imaged at day 0, 3 and at 4 weeks post intra-articular cell injection. Red arrows indicate the site of MSC implantation. Rows of images correspond to the same mouse over time. Control is contralateral knee without surgery or any manipulation of any kind. **B.** Percent of retained void volume compared to day 0, showing MSCs presence in DMM and SHAM operated knee joints up to 4 weeks post-injection via MRI imaging (all ferumoxytol-labeled or unlabeled cells are DiR labeled). **C.** Percent fractional signal loss (i.e. the difference in signal in the void divided by the signal in the surrounding tissue times 100) over time. *p<0.05 between DMM and sham animals at the same time point as determined by Student’s t-test and Tukey post-hoc test. **D.** Persistence of MSCs in mouse joints over a period of 4 weeks, as displayed by DiR fluorescence labeling on all injected MSCs. Mice subjected to DMM+Saline was used as an imaging control.

On the day of cell injection, the average volume of the region of signal loss within the surgical knee did not differ significantly between mice with different surgical preparations with a volume of 0.51±0.38 mm^3^ in DMM mice and 0.39±0.25 mm^3^ in sham mice (Fig 2A). In all DiR+Fe-MSC injected knees the void volume decreased overtime, however, a region of signal loss persisted in all of these knees at the transplant site until end point. A significantly greater percentage of the initial void volume persisted at 3 days and 1-week post DiR+Fe-MSC cell injection in DMM knees compared to sham knees (Fig 2B). By the 4-week end point both surgical groups showed a significant (p<0.0001) reduction in void volume, but DMM surgery mice still retained a void volume of 0.17±0.04 mm^3^ which was significantly greater (p=0.001) volume than the 0.06±0.03 mm^3^ void volume seen in sham surgery mice (Fig 2A).

On the day of cell transplantation, the fractional signal loss (FSL) in DiR+Fe-MSC injected surgical knees did not differ significantly with FSL values of 77.0±6.9 and 75.7±6.3 for DMM and sham knees, respectively (Fig 2C). DMM surgery mice receiving DiR+Fe-MSC did not show a significant reduction in FSL over the time course of the experiment (Fig 2C). Conversely, sham surgery mice showed a significant (p=0.0021) 18% reduction is FSL as early as 3 days post injection. The FSL observed in DMM versus sham surgery mice was significantly different (p≤0.05) at all assessed time points after the day of injection (Fig 2C). By the 4-week end point DiR+Fe-MSC injected DMM and sham surgery knees showed FSL values of 67.6±8.0 and 51.0±9.5, respectively.

On the final day of imaging (at 1, 2 or 4 weeks) both the surgical and contralateral knees were collected and assessed for MSC persistence by fluorescent imaging. DiR fluorescence was detectable in DiR MSC and DiR+Fe-MSC injected DMM surgical knees (Fig 2D). No fluorescence was evident in uninjected contralateral knees nor those injected solely with saline.

In addition to tracking DiR+Fe-MSC implanted cells, longitudinal MRI analysis allowed the evaluation of suspected inflammation in surgical knees over time. DMM surgical knees receiving either vehicle (saline) or DiR+Fe-MSC were scored with values between 1 and 5 where 1 represented no evidence of inflammation and 5 represented severe evidence of inflammation with likely swelling beyond the normal joint space. On the day of cell implantation, the majority of DMM surgical knees showed evidence of inflammation; scores of 4.4±0.7 and 4.0±1.0 were recorded for saline and DiR+Fe-MSC injected DMM knees, respectively (Fig 3A). At the 4-week end point DiR+Fe-MSC injected DMM knees showed significantly less (p=0.01) evidence of inflammation and were scored 1.2±0.4 compared to saline knees scoring 3.6±1.1.

**Figure 3:**
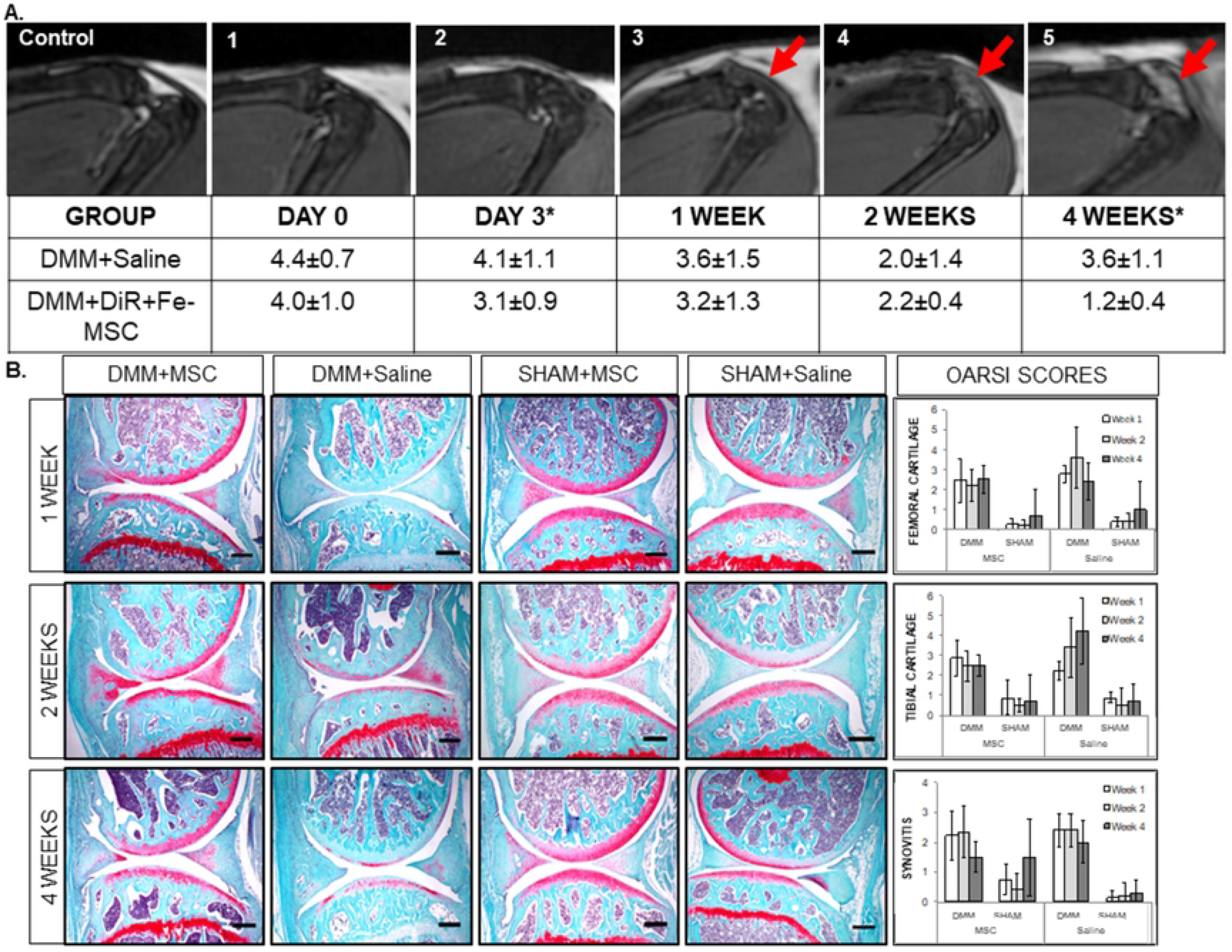
Joint Inflammation, Cartilage Degradation and Synovitis Assessments on DMM or SHAM-operated joints treated with MSCs. **A.** MRI assessment of joint inflammation with representative images to indicate scale. The following scale corresponds to amount of swelling observed in the joint. 1 = Not evident; 2 = Unlikely; 3 = Uncertain; 4 = Likely; 5 = Clear Evidence. The table indicates representative joint inflammation scores on DMM-operated mice treated with or without DiR+Fe-MSC. Significance was found at Day 3 and Week 4 mouse groups (*p<0.05). These images are representative of both DiR+Fe-MSCs and DiR MSCs. **B.** OARSI scoring for of DMM-operated or SHAM-operated joints treated with MSC or saline were assessed on Safranin O-stained sections. No significant difference in cartilage degradation was observed between all groups. The OARSI scores were calculated by including the scores of both DiR+Fe-MSCs and DiR MSCs injection in the DMM surgery groups while only DiR+Fe-MSCs was used in the SHAM-operated joints. Scalebar = 500 μm)

### Retention of iron signal in MSCs

Histological sections were stained against Prussian Blue to detect presence of ferumoxytol in injected MSCs [19]. Prussian Blue staining was found to localize within the synovial joint, indicating that ferumoxytol-labeled MSCs persisted in the joint space up to 4 weeks after intra-articular injection (Fig 4A). Notably, positively stained Prussian-blue labeled MSCs (blue) were detected within macrophages (stained with F480, purple) as early as 1 week after intra-articular injection, suggesting that macrophages engulfed apoptotic MSC bodies during the early inflammatory phase of osteoarthritis (Fig 4B).

**Figure 4:**
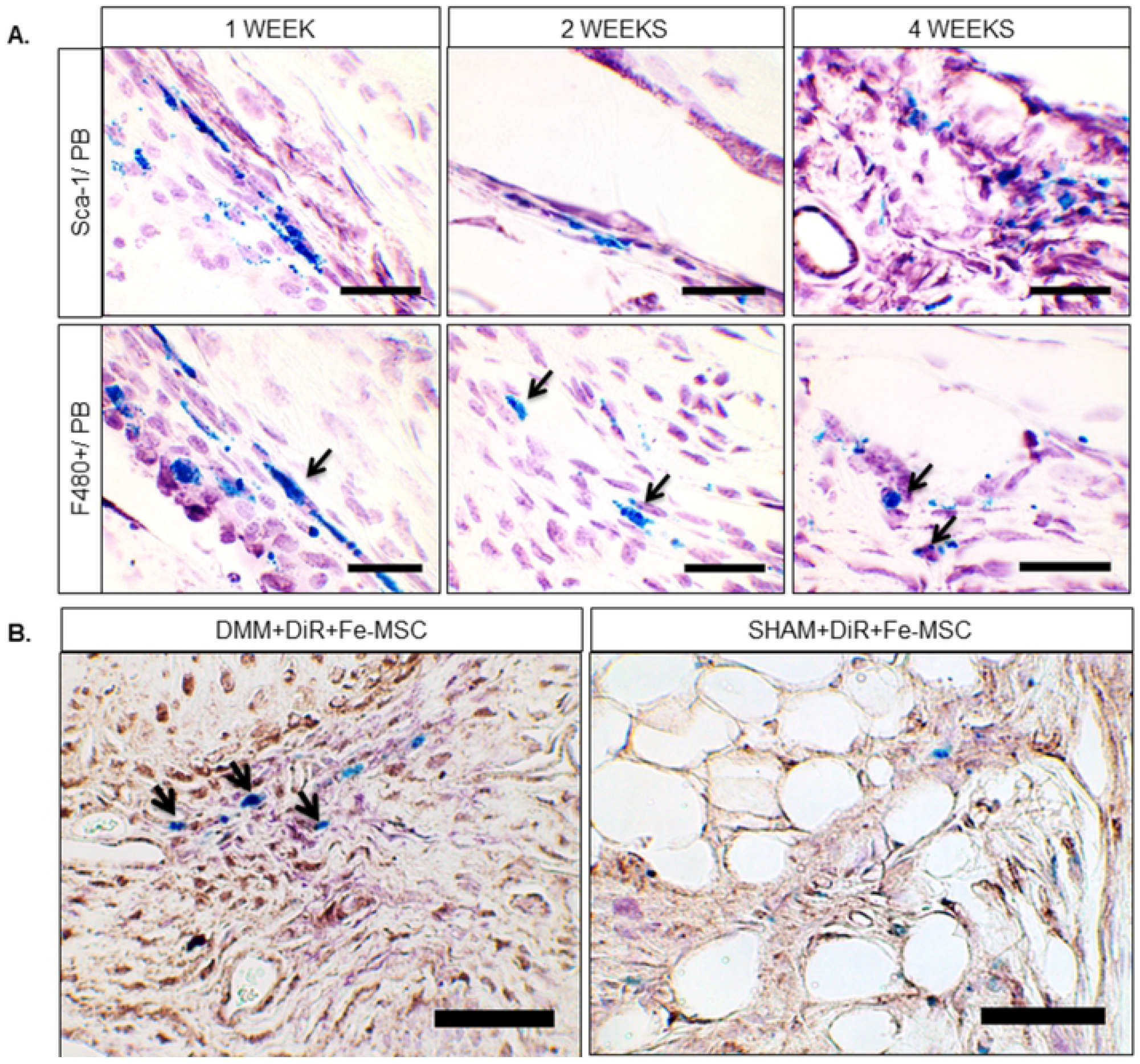
Histopathological Analyses of MSCs in DMM-operated mice at 1, 2, and 4 weeks post-injection. **A.** Mouse knee sections were co-stained with Sca-1 (Purple) and Prussian Blue (PB, blue) to indicate persistence of DiR+Fe-MSC at 1, 2 and 4 weeks post injection. Co-staining of F480 (purple) and Prussian Blue (blue) demonstrated some co-localization of DiR+Fe-MSC cells with macrophages. (Black Arrows.) Scalebar = 50 μm. **B.** Mouse knee sections were triple-labeled with Sca-1 (Purple), Cleaved Caspase-3 (brown), and Prussian Blue (blue), to indicate presence of apoptotic DiR+Fe-MSCs in synovium. Scalebar = 50 μm.

### Histopathology of Synovial Joints

Cartilage degeneration of mouse joints was graded using the OARSI scoring system. No significant changes were found in cartilage degradation between MSC-treated and saline-treated OA-mice. (Fig 3B).

Using the OARSI scoring system for synovial inflammation [18], average synovial inflammation scores (0-4) were determined in DMM-operated mice treated with saline or DiR+Fe-MSCs. Differences in synovial inflammation were detected during the early phase of synovial inflammation (i.e. 3 days) and late inflammation (i.e. 4 weeks) after MSC treatment (p<0.05 respectively).

The substantial decrease in synovial inflammation at 4 weeks after MSC injection corresponded with the prolonged persistence of DiR+Fe-MSCs within the synovial joint, and an increase of homeostatic macrophages at 4 weeks after injection, suggesting that the MSC treatment may be contributing significantly to the infiltration and polarization of macrophages into a more homeostatic phenotype to resolve synovial inflammation within the joint.

### Immunohistochemistry

Medial knee sections were stained against CD206 and F480 to determine the amount of homeostatic macrophage and total macrophage infiltration after MSC injection in OA-induced mice (Fig 5A). We observed that MSC-treatment in DMM mice induced an increasing trend in total macrophage infiltration (p<0.08) compared to saline-treated DMM mice and sham operated mice treated with either MSCs or saline. More notably, there was a significant increase in CD206+ cells in the synovium after MSC treatment in DMM-operated mice, suggesting that MSC treatment led to an increase in macrophage infiltration and polarization towards a homeostatic phenotype (p<0.05, Fig 5B).

**Figure 5:**
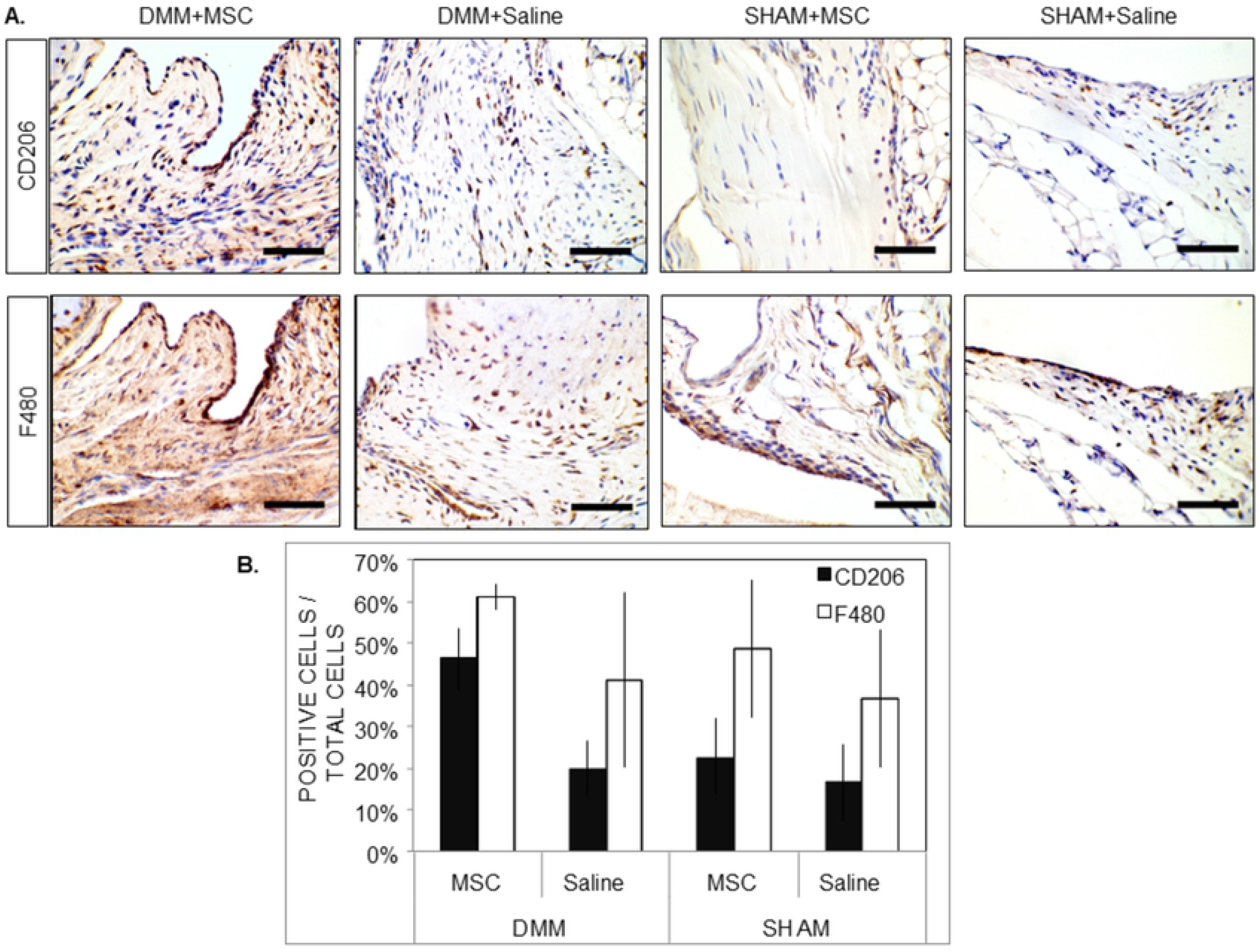
MSC treatment promoted increased proportions of homeostatic macrophages into osteoarthritic synovium at 4 weeks post-treatment. **A.** Sections from DMM-operated and SHAM-operated mice treated with DiR+Fe-MSC or saline were stained with CD206 (brown), F480 (brown), and hematoxylin (blue) to detect homeostatic macrophages, total macrophages and total cells respectively (Scale bar = 50 μm). These images are representative of animals from both DiR+Fe-MSCs and DiR MSC groups. **B.** Quantification of positively stained CD206 or F480 cells. DMM-operated mice treated with MSC yielded significantly more CD206+ cells (homeostatic macrophage) into the synovium than vehicle treated OA mice (*p<0.05). The quantification was done by including both DiR+Fe-MSCs and DiR MSCs injection in the DMM surgery groups while only DiR+Fe-MSCs was used in the SHAM-operated joints.

*Safety of* ferumoxytol-labeled *MSCs vs. unlabeled MSCs* - Safety of ferumoxytol-labeled MSC over a 4-week period was also verified in 35 mice with histology on target organs, and CBC and blood biochemistry (Table S2 and S3). Our data shows no difference in readouts between ferumoxytol-labeled and unlabeled MSCs, saline controls or sham animals.

**Table S2: Blood chemistry results in mice receiving DiR+FeMSCs vs. DiR MSCs at 2 and 4 weeks.**

**Table S3: Pathology report after injection of DiR+Fe-MSCs vs. DiR MSCs at 2 and 4 weeks.** The report shows that Fe-MSCs are safe as assessed by gross pathology of heart and spleen. n/s: No significant findings; H*: The majority of the myocardium appears normal. There is one region of endocardium that has a small amount of fibrin deposition. Duration: subacute; Distribution: focal; Severity: moderate; S*: There are a few areas of decreased density in the periphery of the red pulp. The marginal zones also appear moderately decreased.

## Discussion

Using a non-invasive clinically-approved tracking method, we have for the first time imaged ferumoxytol-labeled MSC persistence in an *immunocompetent clinically translatable* mouse model of surgically-induced osteoarthritis (OA). We showed persistence of MSCs out to four weeks after injection, and importantly illustrated a mechanism of actions involving MSCs polarization of joint-specific macrophages to more homeostatic lineages.

Previously, serial scans of xenotransplanted human MSCs (hMSCs) in an osteochondral rat defect model showed that the iron signal of apoptotic, nanoparticle-labeled hMSCs engulfed by macrophages disappeared faster compared to viable hMSCs in T2-weighted MRI images [20]. This corresponded to poor cartilage repair outcomes in rats receiving apoptotic hMSCs.

Delling et al. similarly used a non-vital ovine MSC group in 12 sheep that had OA-induced by bilateral menisectomy and reported significant differences in hypointensity signaling up to 12 weeks between the vital and non-vital MSC groups [21]. Importantly, they observed reduced Prussian Blue staining in sheep receiving the non-vital MSCs by immunohistology compared to vital MSC groups. They did report increased chondrogenic differentiation surrounding regions of iron nanoparticle detection.

We report for the first time, that Prussian Blue staining by immunohistochemistry is largely visible in cells that co-stain for Sca-1, a marker of bone marrow-derived mesenchymal stromal cells [22]. Interestingly, we also show Prussian Blue co-staining with F480 positive cells, suggestive of iron nanoparticle signal uptake by macrophages in the joint. This likely suggests phagocytosis of apoptotic MSCs by joint macrophages. However, unlike Daldrup et al., we do not see a diminishment in repair properties due to apoptotic MSCs. In fact, the DMM mice showed the same degree of cartilage degradation with or without MSC injection, 4 weeks post-MSC injection. Recently exosomes from pluripotent stem cell-derived MSCs were reported to improve cartilage degradation in a DMM model [23].

However, the authors used repeated MSC exosome injections (every 3 days over a 4-week period) to achieve this effect. Here, we used a single MSC injection, which also decreased the amount of injury to the synovial joint upon delivery of treatment. In addition, there were xeno- and tissue-source differences: Wang et al. used human embryonic stem cell-derived MSCs while we used murine femur-derived MSCs [23].

In fact, we showed that despite the potential apoptosis of MSCs, there was an increased skewing of macrophages to more homeostatic lineages, based on expression of CD206, relative to mice receiving placebo controls. CD206 is a classical marker of more homeostatic macrophages, as a mannose receptor involved in phagocytosis ([24–26]. Recently it has been suggested that apoptosis of MSCs may in fact enhance their immunosuppressive properties, eliciting an IL10, IDO and TGFβ1 suppressive response similar to efferocytosis of apoptotic cell debris [27,28] as we have observed.

Macrophages are key mediatory cells of synovial inflammation or synovitis, a hallmark characteristic of OA pathology [29]. We have recently shown that prevalence of pro-inflammatory monocytes/macrophages in knee OA patients inversely correlates to worsening patient-reported outcomes [14]. Others have similarly reported that prevalence of folate-receptor (i.e., activated) macrophages correlates with OA severity [30]. Pro-inflammatory monocytes/macrophages produce inflammatory and degradatory mediators [31], activate T cells and the complement system [31], and inhibit chondrogenesis in vitro [32].

Others have also previously reported that MSC injection into DMM models does not result in improvement to cartilage scores, consistent with our results in this study, and indeed with our clinical observations in a 12 patient MSC knee OA trial (Chahal et al., in press [7]). Schelbergen et al. have previously shown that adipose tissue-derived (AT) MSCs improved synovitis outcomes, which are pronounced in a collagenase-induced OA (CIOA) model, but not in a DMM model [33]. Unlike this study, which primarily reported AT-MSC-mediated reduced synovitis in the CIOA, we did investigate changes to inflammation even in this mild-inflammatory model, and report that bone marrow MSC treatment does rescue synovial inflammation.

There is a need to specifically track MSCs following infusion to evaluate their migration within the arthritis joint, and to quantify their accumulation and persistence at the target, and elsewhere in the body. This in turn will enable better dosing and design of Phase II clinical investigations. Our findings show feasibility of using ferumoxytol (Feraheme™) for tracking short-term persistence and biodistribution of MSCs. Ferumoxytol as an iron replacement product indicated for the treatment of iron deficiency anemia in adult patients with chronic kidney disease (CKD) (DIN # 02377217). Importantly, we showed no differences in murine and human (Fig S2) MSCs in terms of their viability, cell surface markers or tri-lineage differentiation capacity. Fe-MSCs were similar to unlabeled MSCs in terms of blood chemistry and gross histopathology of target organs (heart and spleen) (Table S2 and S3). This is important to note as some adverse findings have been reported with intravenous use of ferumoxytol at doses that are 600-fold higher than what we propose for ex-vivo labeling of MSCs (only 10% of MSCs will be labeled in a clinical infusion scenario). Recently, off-label use of ferumoxytol as a contrast agent (single 5 mg/kg or 4x doses) in 49 pediatric patients and 19 adults showed no serious adverse events and 4 mild, self-resolving adverse events[34]. This dose is 6.67-fold greater than the 0.75 mg dose we are proposing. Three investigation new drug applications (IND) (NCT02006108, NCT01336803 and NCT02893293) using ferumoxytol as a contrast agent were approved by the FDA; one clinical trial application (CTA, NCT02466828) using ferumoxytol as a contrast agent was approved by Health Canada. These proposed doses are 120-fold lower than the approved intra-venous dose, and about 6.67-fold higher than what we would propose to use in a clinical tracking study of MSCs. In this trial, we plan to inject up to 10% ferumoxytol-labeled MSCs of the total dose (10% labeled MSCs + 90% unlabeled MSCs) and track them at different time points by MRI imaging. Our findings lay the ground work for a clinical pharmacokinetics/pharmacodynamics study of MSCs in osteoarthritis. Our findings confirm an anti-inflammatory mechanism of action of MSCs in osteoarthritis.

## Summary

We have shown that ferumoxytol-labeled murine MSCs can be tracked by MRI imaging in murine osteoarthritic joints without adverse events for up to 4 weeks. Ferumoxytol labeling of MSCs caused no alterations in viability, gene expression (except for HGF) or tri-lineage differentiation potential. Ferumoxytol-labeled MSCs persisted for a longer period of time in surgically-injured DMM mice joints than in SHAM controls. Persistence of the ferumoxytol label in MSCs was confirmed by double staining of Sca-1+ positive cells with Prussian Blue. The presence of Prussian Blue staining in neighboring macrophages argues for efferocytosis of apoptotic MSCs, which has recently been shown as a viable mechanism that enhances MSC immunomodulation. We report for the first time that DMM mice receiving MSCs had an increased proportion of homeostatic polarized macrophages in the arthritic joint than control groups, suggesting that MSCs can have anti-inflammatory effects on OA synovitis. Taken together, our imaging study provides new insights into MSC mechanism of action and lays the groundwork for clinical tracking studies using ferumoxytol-labeled MSCs.

## Acknowledgements

This work has been funded by Stem Cell Network (Clinical Trial Impact grant # 8) as well as the Ontario Institute for Regenerative Medicine (Accelerator grant). Wing-Yee Cheung’s salary was funded in part by The Arthritis Society Young Investigator Operating grant (TAS-YIO 15-321) for Dr. Viswanathan. The authors would like to acknowledge the Pathology Core at The Centre for Phenogenomics as well as the Animal Resources Centres of the University Health Network and of the University of Western Ontario for their technical services. We declare that there was no role of the funding source on the design of the study and collection, analysis, and interpretation of data and in writing the manuscript.

## Notes

**Conflicts of Interest:** Sowmya Viswanathan has a regulatory consulting company, which does not conflict in anyway with any work done for this paper. The other authors have no conflicts of interest.

